# Tomato root transformation followed by inoculation with *Ralstonia solanacearum* for straightforward genetic analysis of bacterial wilt disease

**DOI:** 10.1101/646158

**Authors:** Rafael J. L. Morcillo, Achen Zhao, María I. Tamayo-Navarrete, José M. García-Garrido, Alberto P. Macho

**Author notes:** For correspondence: Rafael J. L. Morcillo, Alberto P. Macho.

## Abstract

*Ralstonia solanacearum* is a devastating soil borne vascular pathogen that is able to infect a large range of plant species, causing an important threat to agriculture. However, the *Ralstonia* model is considerably under-explored in comparison to other models involving bacterial plant pathogens, such as *Pseudomonas syringae* in *Arabidopsis*. Research targeted to understanding the interaction between *Ralstonia* and crop plants is essential to develop sustainable solutions to fight against bacterial wilt disease, but is currently hindered by the lack of straightforward experimental assays to characterize the different components of the interaction in native host plants. In this scenario, we have developed an easy method to perform genetic analysis of *Ralstonia* infection of tomato, a natural host of *Ralstonia*. This method is based on *Agrobacterium rhizogenes*-mediated transformation of tomato roots, followed by *Ralstonia* soil-drenching inoculation of the resulting plants, containing transformed roots expressing the construct of interest. The versatility of the root transformation assay allows performing either gene overexpression or gene silencing mediated by RNAi. As a proof of concept, we used this method to show that RNAi-mediated silencing of *SlCESA6* of tomato roots conferred resistance to *Ralstonia*. Here, we describe this method in detail, enabling genetic approaches to understand bacterial wilt disease in a relative short time and with small requirements of equipment and plant growth space.

**SUMMARY:** A versatile method for tomato root transformation followed by inoculation with *Ralstonia solanacearum* to perform straightforward genetic analysis for the study of bacterial wilt disease.

## INTRODUCTION

*Ralstonia solanacearum*, the causal agent of bacterial wilt disease, is a devastating soil borne vascular pathogen, which shows a world distribution, and is able to infect a large range of plant species, including potato, tomato, tobacco, banana, pepper and eggplant, among others (Jiang et al., 2017; Mansfield et al., 2012). Yield losses caused by *Ralstonia* can rich 80-90% of production in tomato, potato or banana, depending on cultivar, climate, soil and other factors (Elphinstone, 2005). However, the *Ralstonia* model is considerably under-explored in comparison to other models involving bacterial plant pathogens, such as *Pseudomonas syringae* or *Xhantomonas spp*. Additionally, most studies in plant-microbe interactions are focused on the model plant *Arabidopsis thaliana*. Although research using these models has largely contributed to our understanding of plant-bacteria interactions, they do not address the current necessity to understand these interactions in crop plants. Research targeted to understanding the interaction between *Ralstonia* and crop plants is essential to develop sustainable solutions to fight against bacterial wilt disease, but is currently hindered by the lack of straightforward experimental assays to characterize the different components of the interaction. Particularly, tomato, a natural host for *Ralstonia*, is the second most important vegetable crop worldwide and is affected by a plethora of diseases (Jones et al., 1991), including bacterial wilt disease. In this work, we have developed an easy method to perform genetic analysis of *Ralstonia* infection of tomato. This method is based on *Agrobacterium rhizogenes*-mediated transformation of tomato roots (Ho-Plágaro et al., 2018), followed by *Ralstonia* soil-drenching inoculation of the resulting plants, containing transformed roots expressing the construct of interest. The versatility of the root transformation assay allows performing either gene overexpression or gene silencing mediated by RNAi.

Although plant resistance against *Ralstonia* is not well understood, several reports have associated cell wall alterations to enhanced resistance to bacterial wilt (Hernandez-Blanco et al., 2007; Denance et al, 2013). It has been suggested that these cell wall alterations affect vascular development, an essential aspect for the lifestyle of *Ralstonia* inside the plant (Digonnet et a., 2012). Mutations in genes encoding the cellulose synthases *CESA4, CESA7* and *CESA8* in *Arabidopsis thaliana*, have been shown to impair secondary cell wall integrity, causing enhanced resistance to *Ralstonia*, which appears to be linked to ABA signalling (Hernandez-Blanco et al., 2007). Therefore, as a proof of concept for our method, we performed RNAi-mediated gene silencing of *SlCESA6* (*Solyc02g072240*), a secondary cell-wall cellulose synthase, and ortholog of *AtCESA8* (*At4g18780*). Subsequent soil-drenching inoculation with *Ralstonia* showed that silencing *SlCESA6* conferred resistance to bacterial wilt symptoms, suggesting that cell wall-mediated resistance to *Ralstonia* is likely conserved in tomato, and validating our method to carry out genetic analysis of bacterial wilt resistance in tomato roots. Here, we describe this method in detail, enabling genetic approaches to understand bacterial wilt disease in a relative short time and with small requirements of equipment and plant growth space.

## PROTOCOL

Note: important parts of this method involve handling of plant materials *in vitro*, and therefore it is important to keep sterile conditions during all these procedures, including the visualization of *DsRed* fluorescence.

Note II: during all the transformation process, tomato seedlings grow at 25-28 °C and 16/8h light/dark (130 µmol photons m^-2^sec^-1^ light). Plates are sealed with *micropore* tape in order to facilitate gas exchange and transpiration.

### 1. Preparation of Tomato Plants and *Agrobacterium rhizogenes*

1.1. Sterilize tomato seeds (*Solanum lycopersicum* cv. Moneymaker) with 5% (v/v) sodium hypochlorite during 5 minutes. Wash 4-5 times with distilled sterile water and keep the seeds shaking slowly in sterile water over-night to facilitate germination.

1.2. Transfer the tomato seeds to half-strength Murashige and Skoog (1/2 MS) medium without sugars (2.21 g/L MS, 8% w/v agar). Keep the seeds in the dark at 25-28 °C for three days. (Figure 1.A).

**Figure 1.**
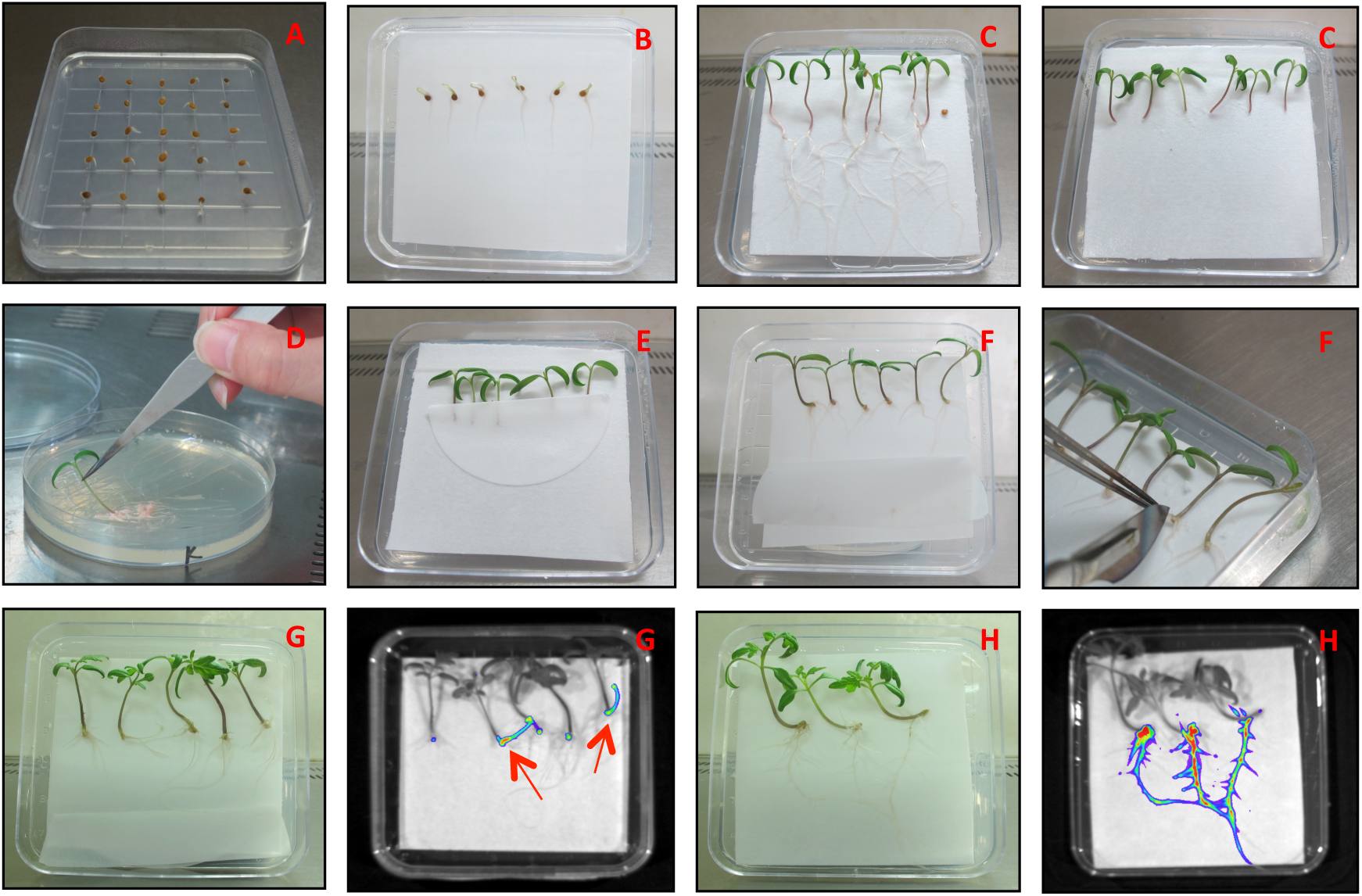
Plant preparation, transformation and selection. **A**.Germination of tomato seeds in half-strength MS (1/2) medium. **B**.Germinated tomato seeds disposed on Whatman paper lying on a plate containing 1/2 MS medium. **C.** Tomato seedlings before and after cutting the radicle and the bottom of the hypocotyl. **D.** Tomato seedling transformation by dipping in bacterial biomass. **E.** Transformed tomato seedlings covered with filter paper to maintain humidity. **F.** Emergence and removal of new (non-transformed) hairy roots. **G.** Selection of transformed tomato hairy roots by visualization of DsRed fluorescence. **H.** Development of the transformed roots as the main roots.

1.3. Autoclave 8.5 cm^2^ *square* filter papers (Whatman paper is ideal; Wahtman International Ltd. Maldstone, England). Place the square filter papers inside 9 cm^2^ square petri dishes containing 1/2 MS medium (place the paper on top of the agar) and dispose six germinated tomato seeds on each plate (Figure 1.B). Seal the plate with *micropore* and incubate the germinated seeds at 25-28 °C during 3-4 days.

1.4. Grow *Agrobacterium rhizogenes* MSU440 in solid LB medium (with appropriate antibiotics) at 28°C two days before plant transformation. In the experiment described in this article, we used *A. rhizogenes* containing pK7GWIWG2_II-RedRoot::CESA6 and empty vector as controls). pK7GWIWG2_II-RedRoot contain a reporter gene, *DsRed*, driven by the *Arabidopsis thaliana* ubiquitin promoter (*pAtUBQ10*), and confers resistance to spectinomycin (50 µg/ml). It is possible to use other vectors for gene overexpression, as reported before (Ho-Plágaro et al., 2018).

### 2. Plant transformation and selection

2.1. Using a sterile scalpel, cut the radicle and the bottom of the hypocotyl of tomato seedlings (Figure 1.C).

2.2.Harvest *A. rhizogenes* biomass from the surface of the LB medium using plastic tips or a scalpel blade and carefully dip the cut tomato seedlings in the bacterial biomass (Figure 1.D).

2.3. After inoculation with *A. rhizogenes*, cover the tomato seedlings with a 2 cm × 4 cm *semi-circular* filter paper, in order to keep a high humidity and facilitate survival and new root development (Figure 1.E).

2.4. Store the transformed tomato seedlings for six-seven days. Then, use a sterile scalpel to cut the new emerging hairy roots (at this step, the roots are not transformed yet) and let the seedlings produce new hairy roots (Figure 1.F).

2.5. Once the *second generation* of new hairy roots appear, remove the filter paper on top of the seedlings, and seal the plate with *micropore* tape again. At this point, the presence of the DsRed fluorescent marker in our transformation vector allows us to visualize the transformation efficiency in the new roots.

2.6. To visualize DsRed fluorescence, it is possible to use a stereomicroscope or any other equipment for plant *in-vivo* imaging. In this work we used the *In Vivo* Plant Imaging System NightShade LB 985 (Berthold Technologies). Mark the positive (red fluorescence) transformed roots and remove the negative non-transformed roots (no red fluorescence) using a sterile scalpel (Figure 1.G).

2.7. Transfer the seedlings showing red fluorescence to a new plate containing 1/2 MS medium, in order to facilitate the development of the transformed root as the main root. Keep the seedlings that do not show red fluorescence in the same plate to check the appearance of fluorescent roots in later time points. Cover the seedlings with a 2 cm x 4 cm *semi-circle* filter paper, seal the plate and incubate the seedlings to let them develop new hairy roots (Figure 1.H).

Note: It is not necessary to eliminate the *A. rhizogenes* using antibiotics. The whatmann paper on top of the MS medium and the lack of sucrose inhibit the spread of the *A. rhizogenes* onto the plate (see Ho-Plagaro et al., 2018).

2.8. Five to seven days after the *first root selection*, repeat the selection process to select new transformed roots and remove non-transformed roots. Transfer the seedlings containing transformed roots to a new plate containing 1/2 MS medium.

### ** Alternative method based on antibiotic selection

A potential limitation of this method consists on the residual growth of non-transformed roots. This is particularly important in the cases where the plasmid used lacks a reporter gene that allows the selection of transformed roots. To solve this problem, we have developed an alternative method based on antibiotic selection, which inhibits the growth of non-transformed roots while allowing the growth of healthy antibiotic-resistant transformed roots. Since *A. rhizogenes* does not induce the transformation of shoots, they are susceptible to the antibiotic, and, therefore, they should be kept separated from the antibiotic-containing medium.

*Alternative 2.5.* Prepare *half-filled* 9 cm^2^ square plates containing 1/2 MS medium with the appropriated antibiotic. Incline the plates approximately 5 degrees during the preparation process to generate an empty space without medium (Figure 2.A), allowing the shoots to grow avoiding contact with the antibiotic.

**Figure 2.**
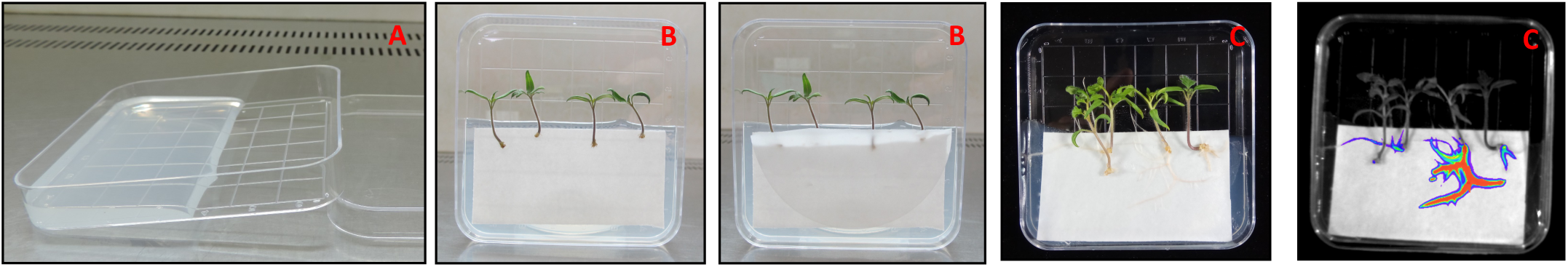
Alternative method based on antibiotic selection. **A.** *Half-filled* square plate containing 1/2 MS medium. **B.** Cut seedlings disposed on Whatman paper lying on 1/2 MS medium with antibiotic, after deletion of first non-transformed hairy roots. **C.** Tomato transformed roots grown on 1/2 MS medium with antibiotic. Seedlings transformed with pK7GWIWG2_II-RedRoot (Kanamycin 50 µg/mL), expressing DsRed, as positive control.

*Alternative 2.6*. After cutting the first non-transformed hairy roots emerged (step 4 in the previous section), transfer the seedlings without roots to the *half-filled* plates using *whatman* and filter papers with the appropriate size (Figure 2.B).

Note: As positive control, it is recommended to transform several seedlings with a plasmid containing the same antibiotic resistance and a reporter gene. In this protocol, we have used pK7GWIWG2_II-RedRoot (Kanamycin 50 µg/mL).

*Alternative 2.7.* Let the seedlings develop new hairy roots and cut those roots that are not in direct contact with the surface of the *wahtman-filter sandwich* papers, since these roots may avoid antibiotic selection. Due to the antibiotic effect, root development may be slower than in plates without antibiotics. New transformed hairy roots appear within 14-18 days after transferring the seedlings without roots to the *half-filled* plates (Alternative 6 step) (Figure 2.C).

### 3. Transfer to inoculation pots

3.1. Prepare inoculation pots where the surface of the roots will be exposed to the bacterial inoculum (Jiffy pots are ideal; Jiffy Products International A.S., Norway): soak the jiffy pots with water, pour off any excess water, and place them in a plastic planting tray. Transfer the selected seedlings with transformed roots to jiffy pots using tweezers (Figure 3.A).

**Figure 3.**
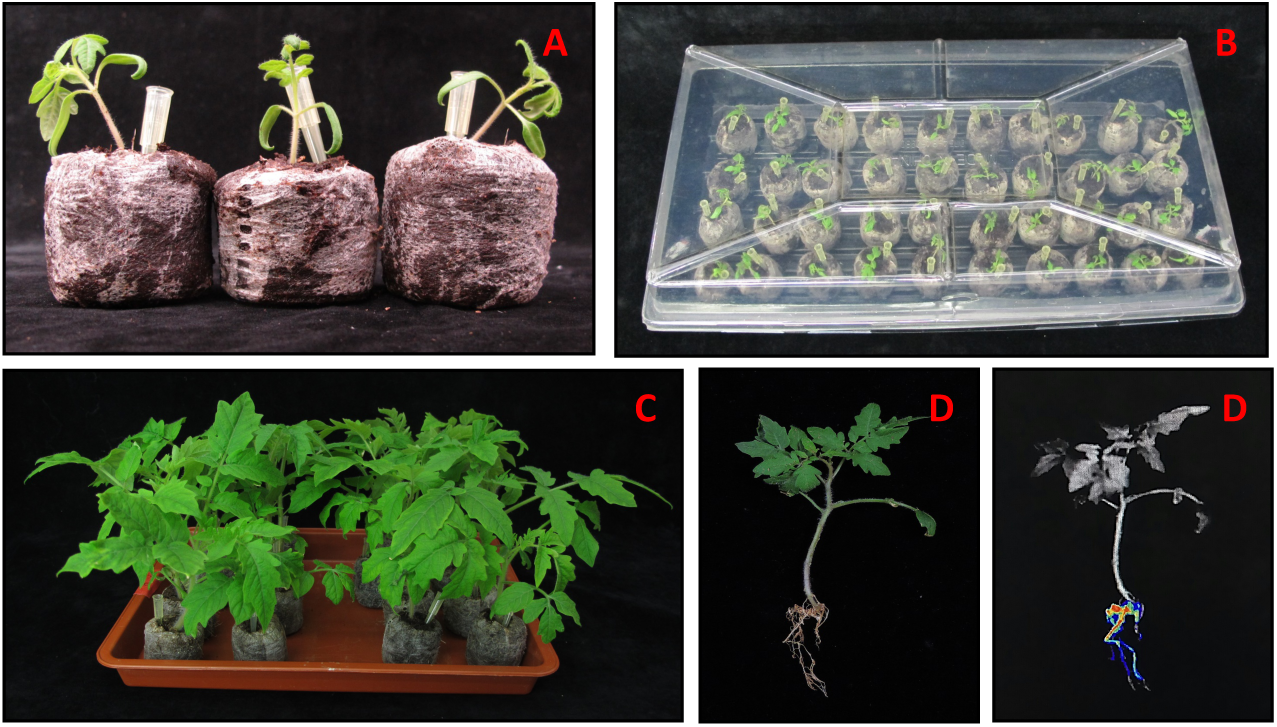
Transference of transformed tomato to inoculation pots. **A.** Root-transformed tomatoes transferred to full-soaked jiffy pots. **B.** Transformed tomatoes in jiffy pots covered with a plastic lid to maintain high level of humidity. **C.** Two-to-three week old root-transformed tomatoes, after transference to jiffy pots, ready to be inoculated. **D.** Red fluorescence in roots of three week-old transformed tomato plants.

3.2. Cover the tray with plastic wrap or transparent lid and keep them at 25-28 °C and 65% humidity (16/8h light/dark; 130 µmol photons m^-2^sec^-1^ light), to maintain high level of humidity (Figure 3.B). Remove the cover after five or six days.

3.3.The transformed tomato plants are ready for inoculation two-to-three weeks after transferring them to soil (Figure 3.C)

Note: we considered the possibility that, during the growth on soil, tomato plants may produce new roots that are not transformed. To determine the extent of this phenomenon, we visualized red fluorescence in roots of three week-old transformed tomato plants. Most roots (80%-100%) showed red fluorescence, indicating that they are indeed transformed (Figure 3.D).

### 4. Soil-drenching inoculation

4.1. Grow *Ralstonia solanacearum* (we used the reference *R. solancearum* strain GMI1000 in this protocol) in phi liquid medium (Table I; Sang et al., 2018) in an orbital shaker (200 rpm) at 28 °C until stationary phase.

4.2. Determine bacterial numbers by measuring the optical density of the bacterial culture at 600 nm (OD_600_). Dilute the bacterial culture with water to an OD_600_ of 0.1 (in our conditions, this corresponds to approximately 10^8^ colony-forming units (cfu)/ml).

4.3. Place 16-20 jiffy pots containing transformed tomato plants in an inoculation tray (29 × 20 cm).

4.4. Pour 300 ml (∼15 mL for plant) of bacterial inoculum (OD_600_ of 0.1) into the tray containing the jiffy pots. Let them soak in the inoculum for 20 minutes (Figure 4.A).

**Figure 4.**
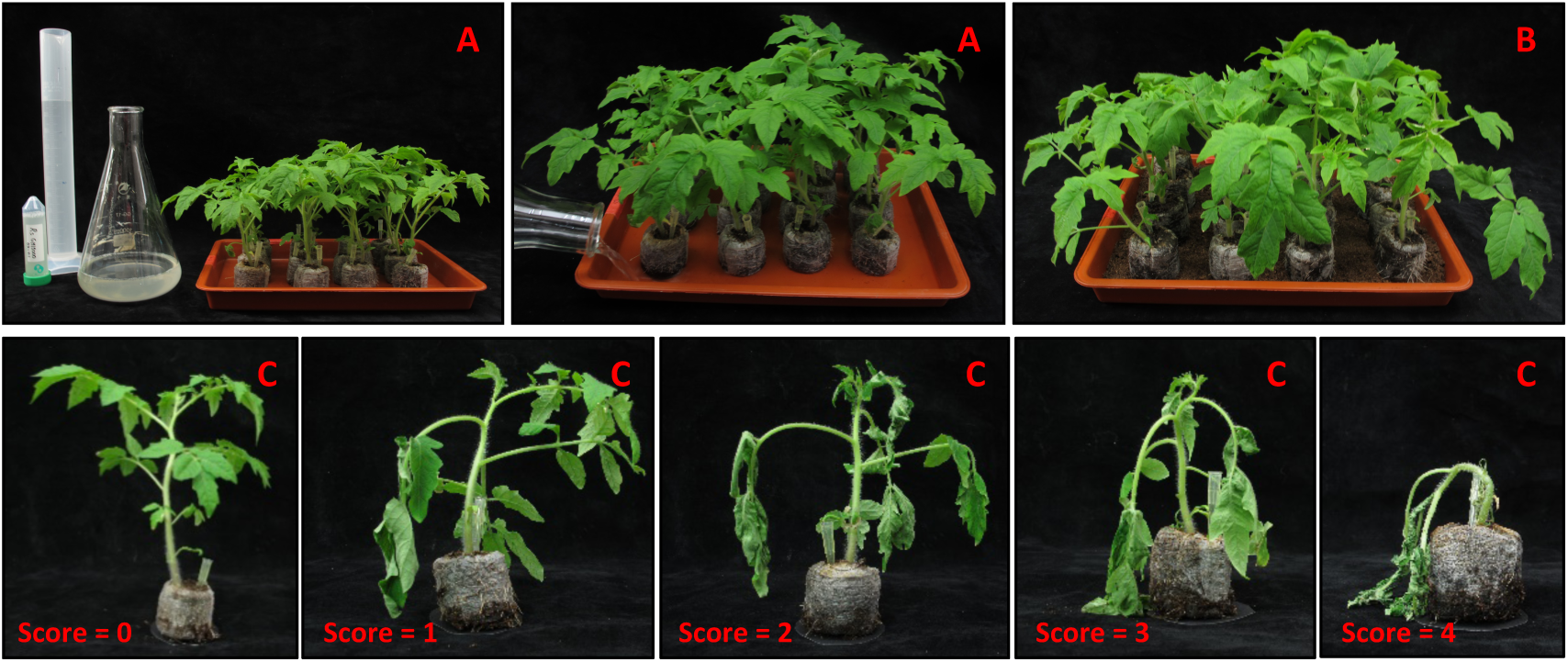
Ralstonia soil-drenching inoculation. **A**.Materials needed for *R. solanacearum* GMI1000 inoculation. Soil-drenching of transformed tomatoes in jiffy pots with *R. solanacearum* GMI1000 inoculum. **B.** Inoculated tomatoes disposed on a layer of potting soil. **C.** *Ralstonia* disease symptoms scale ranging from 0 (no symptoms) to 4 (complete wilting).

4.5. Prepare a new tray with a layer of potting soil (*Pindstrup* substrate). Move the inoculated jiffy pots into the new tray (Figure 4.B), and place the trays in a growth chamber with 75% humidity, 26-28 °C, and a photoperiod of 12 h light and 12 h darkness (130 µmol photons m^-2^sec^-1^ light).

### 5. Determination of infection parameters and statistical analysis

The scoring of disease symptoms can be performed as previously described (Remigi et al., 2011; Wang et al., 2016), using a scale ranging from 0 (no symptoms) to 4 (complete wilting), each day after *Ralstonia* inoculation during two weeks. Score 2 corresponds to 50% of the leaves showing wilting symptoms (Figure 4.C). The disease index data are collected from the same experimental unit (each plant) over time according to an arbitrary scale from 0 to 4 (Figure 4.C), and do not follow a Gaussian distribution, ruling out the use of standard tests for parametric data. Moreover, we have to consider the high variation existing in this kind of experiments, due to the colonization and infection dynamics. Below we considered different methods for the representation and statistical analysis of symptoms associated to *Ralstonia* infection:

5.1. As standard approach, we used a U Mann-Whitney two-tailed non-parametric test to compare both control (EV) and *SlCESA6*-RNAi infection curves. According with this analysis, the medians of both curves appear to be non-significant (*P value* = 0.16).

5.2. We also quantified the Area Under the Disease Progress Curve (AUDPC), which allows combining multiple observations of disease progress into a single value. AUDPC showed a higher value for control (EV; 28.75 ± 4.75) than *SlCESA6*-RNAi (21.01 ± 4.74) inoculated plants at the end of the infection process, indicating an enhanced of *SlCES6*-silenced plants to *Ralstonia* infection (Figure 5.B).

**Figure 5.**
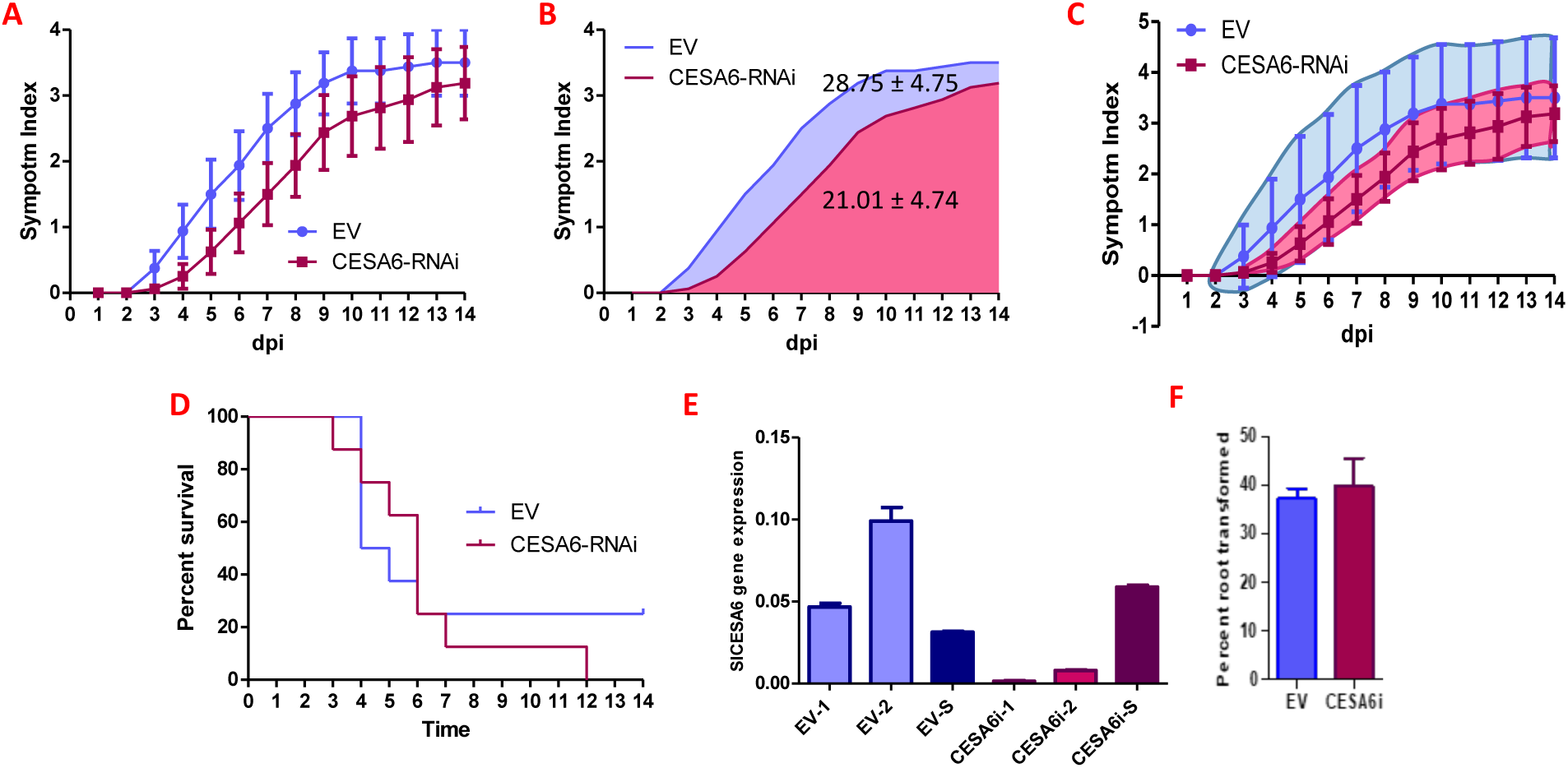
*SlCESA6* silencing enhanced resistance to Ralstonia. **A.** Disease symptoms of RNAi-mediated *SlCESA6*-silenced (CESA6-RNAi) and Empty Vector (EV) root-transformed tomato plants upon inoculation with *Ralstonia solanacearum*. Values correspond to the means±SE of eight plants. **B.** Area under the disease progress curve of plants shown in A. **C.** 95% Confidence Interval of plants shown in A. **D.** Percent of surviving plants shown in A. **E.** Gene expression analysis by qRT-PCR of RNAi-mediated *SlCESA6*-silenced (CESA6i) and Empty Vector (EV) root-transformed tomato plants in roots (1,2,3) and shoot (S). Values correspond to the means ±SE of three technical replicates. **F.** Transformation rate of Empty Vector (EV) and RNAi-mediated *SlCESA6*-silenced (CESA6i) root-transformed tomato plants of two independent experiments. Values correspond to the means ±SE, after two rounds of selection.

5.3. Confidence intervals (CI) offer a way of estimating, with high probability, a range of values in which the population value (or parameter) of a given variable is found. As we show in Figure 5.C, the area of 95% CI for control (EV) and *SlCESA6*-RNAi infection curves estimate a higher chance of resistance when *SlCESA6* is silenced.

5.4. Disease index values can be transformed into binary data, considering a disease index lower than 2 corresponding to “0”, and a disease index equal or higher than 2 corresponding to “1” (Remigi et al., 2011). This allows the representation of a survival curve after *Ralstonia* inoculation. This transformation is based on the observation that, once the plants start developing clear symptoms (disease index of 2), they are considered as “infected” and will die as a consequence of this infection. The differences in the survival rate between EV and *SlCESA6*-RNAi plants were not statistically significant according to a Gehan-Breslow-Wilcoxon Statistical Test (*P value* = 0.13) (Figure 5.D).

Nevertheless, we also advocate to interpret the data based on the biological evidences provide by the consistent phenotype observed among different replicates and not only based on means comparison between samples with a standard *P* value ≤ 0.05, according to the current opinion of interpret data based not only on statistical threshold (Amrhein et al., 2019).

### 6. Gene expression analysis

The expression of the transgene or the silencing of the target gene can be determined by reverse transcription (RT) PCR or by quantitative RT-PCR (qRT-PCR). In this protocol, we always use qRT-PCR to provide a quantitative assessment of gene expression.

6.1. It is recommended to keep several transformed plants without *Ralstonia* inoculation to analyse gene expression. Extract RNA from a representative part of the transformed root system (to evaluate the effect on the target gene) and leaves (as internal control). In this protocol, we used the RNeasy Plant Mini Kit (Qiagen, MD, USA) following the manufacturer’s instructions.

6.2. Synthesize cDNA using 1 µg of total DNase-treated RNA, using the iScript™ cDNA synthesis kit (Bio-Rad).

6.3. Analyse gene expression of the target genes by qRT-PCR. The relative transcription levels were calculated using the 2^-ΔΔCT^ method (Livak andSchmittgen, 2001), using *SlEFα-1* as housekeeping gene. Primer sequences are listed in Table 2.

## REPRESENTATIVE RESULTS

Figure 5 shows the development of disease symptoms of tomato plants with roots transformed with an empty vector, and plants with roots transformed with an RNAi construct targeting *SlCESA6* (*Solyc02g072240*). The results show that silencing *SlCESA6* enhanced resistance disease symptoms upon inoculation with *R. solanacearum* (Figure 5.A).

The expression of *SlCESA6* was analyzed in 3 randomly selected transformed roots before the inoculation step, showing that the enhanced resistance to *Ralstonia* correlates with the reduced expression of *SlCESA6* (Figure 5.C).

Using this method, we typically obtain a transformation rate of 35-40% after two rounds of selection (Figure 5.F); this value can be increased by performing additional selection rounds (Ho-Plagaro et al., 2018).

## DISCUSSION

*Ralstonia solanacearum* poses an important threat to agriculture; however, its interaction with natural hosts of agricultural importance is still poorly understood compared with other bacterial pathogens, especially in crops. In most cases, genetic analysis is hindered by the time and expenses required to genetically modify host plants. To address this problem and facilitate genetic analysis of *R. solanacearum* infection in tomato, we have developed an easy method based on *Agrobacterium rhizogenes*-mediated transformation of tomato roots (Figure 1), followed by soil-drenching inoculation (Figure 3). The transformed roots are selected using a fluorescence reporter gene (*DsRed* in this protocol) (Figure 1). Additionally, we have also developed an alternative method based on antibiotic selection that does not damage the non-transformed aerial part (Figure 2). The transformation protocol is based on that described by Ho-Plagaro et al. (2018) with several modifications. The versatility of this method allows multiple additional assays in transformed roots, such as those to analyse plant physiology, response to chemical treatments, and/or responses to different biotic and abiotic stresses. After transferring the transformed plants to soil, we considered the possibility that plants develop new roots that may not be transformed. We explored this possibility by observing the fluorescence from DsRed in the root system of transformed plants two weeks after transferring them to soil (before *Ralstonia* infection). The results showed red fluorescence in the majority of the roots (Figure 3D).

T-DNA transfer by *Agrobacterium* is integrated in the host genome in a random manner, and, therefore, this method generates a heterogeneous population of tomato plants with different expression level of the target gene. *R. solanacearum* infection normally causes the death of the plants, which, subsequently, hinders the collection of root samples after the experiment to analyse the gene expression of each plant. To analyse the expression level of the target gene during the experiments, we select two to three root-transformed plants before the inoculation step, as representative sample of the population that will be inoculated. The Figure 5 shows that the reduction of disease symptoms in tomato plants correlates with the efficiency of *SlCESA6* silencing before the inoculation step. Therefore, and despite of the limitations of the protocol, our representative results clearly demonstrate that this method is a powerful tool to study candidate genes involved in resistance or susceptibility to *R. solanacearum* in tomato. The example shown in this manuscript allowed us to determine that knocking-down *SlCESA6*, a secondary cell wall-related cellulose synthase, enhances resistance to infection by *R. solanacearum*, resembling previous observations using its ortholog *AtCESA8* (*At4g18780*) from *Arabidopsis thaliana* (Hernandez-Blanco et al., 2007).

## ACKNOWLEDGMENTS

We thank all lab members of the Macho laboratory for helpful discussions, Alvaro Lopez-Garcia for statistical advice, and Xinyu Jian for technical and administrative assistance during this work. We thank the PSC Cell Biology core facility for assistance with fluorescence imaging. This work was supported by the Shanghai Center for Plant Stress Biology (Chinese Academy of Sciences) and the Chinese 1000 Talents program.

## REFERENCES

1 Amrhein, V., Greenland, S. and McShane, B. Retire statistical significance. Nature. 567, 305–307 (2019)

2 Denancé, N., Ranocha, P., Oria, N., Barlet, X., Rivière, M.P., Yadeta, K.A., Hoffmann, L., Perreau, F., Clément, G., Maia-Grondard, A., van den Berg, G.C., Savelli, B., Fournier, S., Aubert, Y., Pelletier, S., Thomma, B.P., Molina, A., Jouanin, L., Marco, Y., Goffner, D. Arabidopsis *wat1 (walls are thin1*)-mediated resistance to the bacterial vascular pathogen, *Ralstonia solanacearum*, is accompanied by cross-regulation of salicylic acid and tryptophan metabolism. Plant J. 73(2), 225–39 (2013).

3 Digonnet, C., Martinez, Y., Denancé, N., Chasseray, M., Dabos, P., Ranocha, P., Marco, Y., Jauneau, A. and Goffner, D. Deciphering the route of *Ralstonia solanacearum* colonization in *Arabidopsis thaliana* roots during a compatible interaction: focus at the plant cell wall. Planta. 236(5), 1419–31 (2012).

4 Elphinstone, J.G. The current bacterial wilt situation: a global overview. In: Allen C, Prior P, Hayward AC, editors. Bacterial Wilt Disease and the Ralstonia solanacearum Species Complex. American Phytopathological Society Press; St Paul, MN. pp. 9–28 (2005).

5 Hernández-Blanco, C., Feng, D.X., Hu, J., Sánchez-Vallet, A., Deslandes, L., Llorente, F., Berrocal-Lobo, M., Keller, H., Barlet, X., Sánchez-Rodríguez, C., Anderson, L.K., Somerville, S., Marco, Y., Molina, A. Impairment of cellulose synthases required for Arabidopsis secondary cell wall formation enhances disease resistance. Plant Cell. 19(3), 890–903 (2007).

6 Ho-Plágaro, T., Huertas, R., Tamayo-Navarrete, M.I., Ocampo, J.A. and García-Garrido, J.M. An improved method for Agrobacterium rhizogenes-mediated transformation of tomato suitable for the study of arbuscular mycorrhizal symbiosis. Plant Methods. 14, 34 (2018).

7 Jiang, G., Wei, Z., Xu, J., Chen, H., Zhang, Y., She, X., Macho, A.P., Ding, W. and Liao B. Bacterial Wilt in China: History, Current Status, and Future Perspectives. Front Plant Sci. 11(8), 1549 (2017).

8 Jones, J.B., Jones, J.P., Stall, R.E., Zitter, T.A. Compendium of Tomato 1094 Diseases. St Paul, MN: APS Press (1991). Mansfield, J., Genin, S., Magori, S., Citovsky, V., Sriariyanum, M., Ronald, P., Dow, M.A.X., Verdier, V., Beer, S.V., Machado, M.A. and Toth, I.A.N. Top 10 plant pathogenic bacteria in molecular plant pathology. Molecular plant pathology. 13(6), 614–629 (2012)

9 Remigi, P., Anisimova, M., Guidot, A., Genin, S. and Peeters, N. Functional diversification of the GALA type III effector family contributes to Ralstonia solanacearum adaptation on different plant hosts. New Phytol. 192, 976–987 (2011).

10 Sang, Y., Wang, Y., Ni, H., Cazalé, A.C., She, Y.M., Peeters, N., Macho AP. The Ralstonia solanacearum type III effector RipAY targets plant redox regulators to suppress immune responses. Mol Plant Pathol. 19(1), 129–142 (2018).

11 Wang, K., Remigi, P., Anisimova, M., Lonjon, F., Kars, I., Kajava, A., Li, C.H., Cheng, C.P., Vailleau, F., Genin, S. and Peeters, N. Functional assignment to positively selected sites in the core type III effector RipG7 from Ralstonia solanacearum. Mol. Plant Pathol. 17, 553–564 (2016).

